# Systematic modulation of sensorimotor learning by domain-specific working memory

**DOI:** 10.1101/2025.04.11.648414

**Authors:** Sean R. O’Bryan, Joshua Liddy, Joo-Hyun Song

**Affiliations:** Department of Cognitive and Psychological Sciences, Brown University, Providence, RI 02912; Department of Kinesiology, University of Massachusetts – Amherst, Amherst, MA 01003; Carney Institute for Brain Science, Brown University, Providence, RI 02912

**Keywords:** Adaptation, Implicit learning, Individual differences, Motor learning, Spatial attention, Working memory

## Abstract

People differ in how quickly they learn and adapt sensorimotor skills, and these differences have been linked to individual variation in working memory capacity (WMc). In tasks that can be supported by cognitive strategies, visuospatial WMc has been proposed as a key contributor. However, it remains unclear whether this association reflects domain-specific mechanisms or domain-general executive resources, and whether it extends beyond spatial memory to other visual features. Here, we systematically tested whether domain-specific or domain-general WMc predicts adaptation across three visuomotor adaptation (VMA) tasks known to differentially engage explicit and implicit learning systems. After obtaining independent measures of spatial and feature-based WMc, healthy subjects completed (1) a standard VMA task, which engages both explicit and implicit learning systems; (2) a delayed feedback VMA task, which isolates explicit learning; and (3) a clamped feedback VMA task, which isolates implicit learning. Our results provide converging evidence in support of a domain-specific association between spatial WMc and individual differences in learning. In Experiments 1 and 2, greater spatial WMc was associated with robust increases in explicit learning whereas featured-based WMc was not associated with learning outcomes. Surprisingly, in Experiment 3, greater spatial WMc was associated with reduced implicit learning, suggesting an interaction between latent cognitive capacities and implicit learning not accounted for by traditional models. These results shed light on the precise cognitive mechanisms underlying sensorimotor adaptation and provide novel insight into domain-specific links between spatial WM and motor learning.

---

Motor skills enable us to successfully navigate and interact with the world around us. Most of the time, we climb stairs without tripping, transport hot coffee without spilling, and navigate busy roadways without much explicit thought. However, when we start to learn new skills, they often feel cognitively demanding. To beginners, for instance, learning to golf or play the guitar may feel impossible at first. Indeed, entire industries have been built around the promise of strategies and training routines that will facilitate effective and efficient learning of such complex skills.

However, despite following the same training routines, there are often considerable individual differences in the speed and fluency with which people learn new skills and adapt existing ones (Anguera et al., 2010; 2012; Christou et al., 2016; de Brouwer et al., 2022; Holland et al., 2019; Seidler, 2006; Standage et al., 2023; Wang et al., 2022). Understanding the nature of this variability requires acknowledging that sensorimotor learning depends not only on skilled movement production, but also on higher-level cognitive processes to flexibly and dynamically generate and refine actions. Given the pervasive influence of cognitive control and explicit strategies in motor skill acquisition (for review, see Seidler et al., 2013; Tsay et al., 2024), investigating how and whether different cognitive capacities support adaptive motor control would significantly advance our understanding of diverse learning outcomes across individuals.

Mounting evidence implicates visual working memory capacity (WMc) as an influential cognitive ability involved in sensorimotor learning (Anguera et al., 2010; 2012; Christou et al., 2016; Holland et al., 2019; McDougle & Taylor, 2019; Seidler, Bo, & Anguera, 2012). Working memory enables the temporary storage, maintenance, and manipulation of information in the mental workspace (Baddeley, 1986; 2003; Cowan, 2001; Logie, 1995; 2003). Although the number of items that can be reliably held in WM is limited (1-4 items; Luck & Vogel, 2013; Vogel & Awh, 2008), WMc varies across individuals in a trait-like manner (Dai et al., 2019; Eng et al., 2005; Nozari & Martin, 2024; Vogel & Machizawa, 2004). Accordingly, WMc predicts success in domains such as mathematics (Beilock & Carr, 2005), reading comprehension (Slattery et al., 2021), academic achievement (Alloway & Alloway, 2010), and navigation (Blacker et al., 2017). Working memory is comprised of functionally distinct modules which operate in a coordinated manner to enable flexible cognition (Baddeley & Hitch, 1974; Baddeley, 2003; Logie, 1995; 2003). While the lateral PFC serves a domain-general role in gating the information stored in WM (Courtney, 2004; Hautzel et al., 2002; Postle et al., 2003; Serences et al., 2004), the contents of WM are held in sensory-specific stores or “buffers” which house representations from different modalities: the phonological loop, which maintains auditory-verbal representations, and the visuospatial sketchpad, which maintains non-verbal visual representations (Baddeley et al., 1999; Repovs & Baddeley, 2006). Relevant to the current study, the visuospatial sketchpad represents visual information of various formats at a more granular, domain-specific level in hierarchically organized sensory regions known to process this information during perception (Jonides et al., 2005; for review, see Serences, 2016; Zimmer, 2008). In particular, spatial information (e.g., locations in egocentric or allocentric space) is maintained in the dorsal extrastriate and parietal cortex (Courtney et al., 1998; Grefkes & Fink, 2005; Jordan et al., 2001; Postle et al., 2000), while non-spatial visual information corresponding to features such color, shape, and texture are maintained in the ventral occipitotemporal cortex (Courtney et al., 1996; Hautzel et al., 2002; Postle et al., 2000; Ungerlieder, Courtney, & Haxby, 1998). Thus, the sites of WM storage in the visual modality can be highly specific, determined by both sensory input and higher-order attentional control processes which filter it.

Whether sensorimotor learning is supported by domain-specific or domain-general visual WM resources is currently unknown. Previous studies have linked independent measures of spatial cognition to motor learning outcomes (Anguera et al., 2010; 2012; Christou et al., 2016; Guo & Song, 2023; Holland et al., 2019), leading to the tacit assumption that spatial WM is exclusively involved. Although these results suggest that certain spatial tasks requiring mental rotation (Anguera et al., 2010; 2012; Guo & Song, 2023; McDougle & Taylor, 2019), passive location-based WM (Christou et al., 2016), or both (Holland et al., 2019) capture between-subject variability relevant to motor learning, they fail to distinguish whether these associations are a driven by domain-general (i.e., executive) or domain-specific (i.e., sensory recruitment) processes. Moreover, the relationship between feature-based WM (more broadly referred to as “object” or “visual” WM) and sensorimotor learning has not been examined. This is significant because domain-specific ventral and dorsal visual pathways play dissociable roles in action planning and execution, where feature-based representations in the ventral stream support the perceptual maintenance of targets in object- or scene-centered coordinates (Goodale & Milner; 1992; Goodale & Westwood, 2004; Goodale et al., 2004). Dual task interference studies suggest that mental rotation depends directly on the storage of discrete visual features and not on the storage of spatial locations in WM (Hyun & Luck, 2007). From this perspective, mental rotation should engage both feature and spatial WM subsystems, with the former holding the representation of an object in mind, and the latter performing the rotation itself (Heil & Rolke, 2002). Finally, it is critical to consider possible confounds unrelated to WM that could drive the observed associations. The presence (or absence) of a relationship between a WM task and learning outcomes could be driven by factors such as visual complexity, task difficulty, or the performance distributions associated with a given dependent measure – possibilities that have not been ruled out. In summary, although multiple studies implicate spatial WM in sensorimotor learning, there is limited evidence that this association is domain-specific; only that it is *modality*-specific.

Our current approach was guided by two primary goals. First, we sought to determine whether the association between visual WMc and sensorimotor learning is, in fact, domain-specific, such that it depends on memory for spatial information and not memory for discrete features. Accordingly, we designed two passive WM tasks with equated response demands and visual properties to more directly index spatial and feature-based representations across participants. Second, we sought to characterize how different types of visual WMc—spatial and feature-based—relate to sensorimotor learning across tasks that engage distinct learning systems. To this end, we compared performance across sensorimotor tasks known to differentially recruit implicit and explicit learning mechanisms, allowing us to assess whether these relationships vary depending on the learning system engaged.

To address these aims, we employed a well-established motor learning paradigm– visuomotor adaptation (VMA)—that allows for the dissocation of implicit and explicit learning processes. In VMA tasks, participants perform reaching movements while receiving altered visual feedback about their hand position (e.g., Krakauer et al., 2000; McDougle et al., 2015; Mazzoni & Krakauer, 2006; Song & Bédard, 2015; Shadmehr et al., 2010; Smith et al., 2006; Taylor & Ivry, 2011; Taylor et al., 2014). This perturbation induces both a sensory prediction error (SPE), a mismatch between expected and actual sensory feedback, and a task error, where the observed outcome deviates from the intended outcome. These distinct error signals engage separate learning mechanisms: implicit adaptation, which unfolds incrementally and is driven by sensory prediction error (Mazzoni & Krakauer, 2006; Morehead et al., 2017; Wolpert et al., 1998; Wolpert & Ghahramani, 2000), and explicit adaptation, a cognitively mediated process that enables rapid, strategic corrections in response to task error (McDougle & Taylor, 2019; Taylor et al., 2010; Taylor, Krakauer, & Ivry, 2014). While implicit adaptation is considered largely automatic (Morehead et al., 2017; Mazzoni & Krakauer, 2006), individual differences in VMA performance have been linked to variability in explicit, WM-dependent strategies (Benson, Anguera, & Seidler, 2011; Christou et al., 2016; Standage et al., 2023; Taylor & Ivry, 2011). However, because most VMA tasks engage both explicit and implicit processes, the extent to which their relationship with WM generalizes across dissociable learning contexts remains an open question.

In three samples of healthy adults, we administered spatial and feature-based working memory tasks followed by VMA tasks designed to tax explicit learning, implicit learning, or both. If visual WMc contributes to adaptation through domain-general executive processes, then both spatial and feature-based WMc should similarly predict learning outcomes. Alternatively, if WMc supports adaptation via domain-specific spatial mechanisms, then spatial—but not feature-based— WMc should be selectively predictive of learning outcomes. In Experiment 1, we found domain-specific effects of visual WMc in an abrupt VMA task with continuous feedback: individuals with higher spatial WMc showed enhanced early adaptation. In Experiment 2, we test whether this association generalized to a context where implicit learning was disrupted by delaying visual feedback (Brunder et al., 2016). Spatial WMc again predicted individual differences in adaptation not only during early learning, but across the entire learning bout. In Experiment 3, we used a clamped feedback paradigm (Morehead et al., 2017) to isolate implicit adaptation and observed that higher spatial WMc suppressed implicit adaptation, suggesting that cognitive capacities can modulate even putatively automatic learning processes. These findings reveal a surprising and consistent pattern: spatial WMc—but not feature-based WMc—tracks adaptation across multiple learning contexts. Our results underscore the importance of distinguishing between subsystems of visual WM in understanding individual differences in sensorimotor learning.

## Results

### Spatial and feature-based working memory capacity

To obtain individual estimates of WMc in dissociable visual domains, participants from all three experiments first completed working memory tasks which probed memory for either spatial locations or visual features (Fig. 1A). The spatial and feature-based tasks were visually identical and differed only in the dimension participants were instructed to attend—dot location or color. Memory load varied randomly from trial-to-trial, with each display containing either three or five colored dots during a 400 ms encoding period. After a 2500 ms delay, participants were shown a single colored dot and indicated via keypress whether it matched any of the previously presented dots on the task-relevant dimension (location or color or location). Each participant completed two blocks of each task, administered in pseudorandom order.

**Figure 1.**
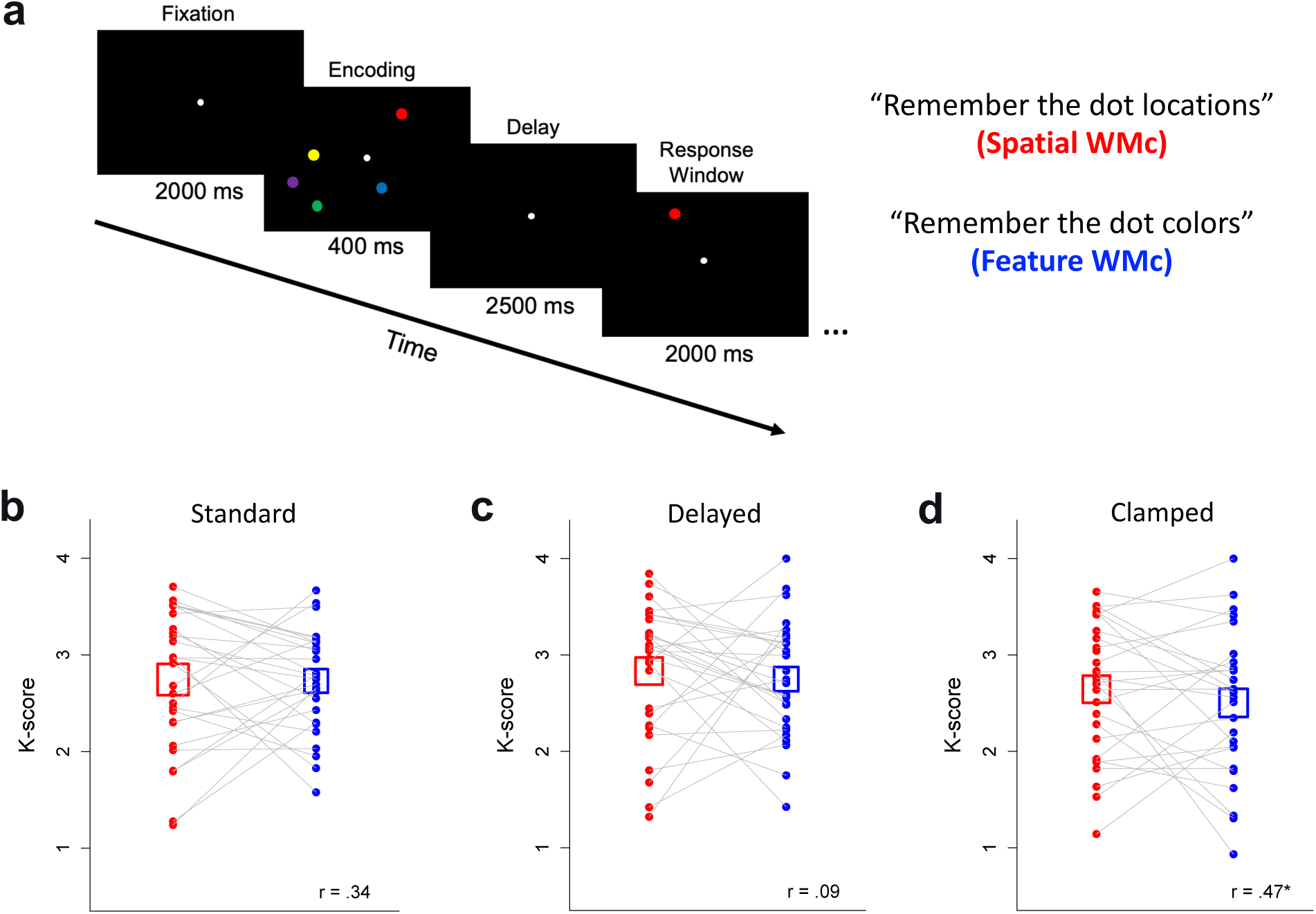
Estimating spatial and feature-based working memory capacity. A) Example trial sequence for the working memory task. Participants were instructed to memorize either the locations (spatial) or colors (feature) of 3 - 5 dots presented during the encoding period. Following a delay, participants responded with a keypress corresponding to whether a single dot presented during the retrieval period was a match or non-match on the attended dimension with any dot presented in the encoding display. The pictured sequence depicts a non-match on the spatial dimension and a match on the feautre dimension for a 5-item array. B-D) Capactiy estimates (*K*-scores) for the spatial (red) and feature (blue) tasks in Exps. 1-3. Squares depict mean *K*-scores +/- SEM. Points depict *K*-scores for individual subjects, where gray connecting lines reflect differences in *K* between the spatial and feature tasks for each subject. * = *p* < 0.05 for the correlation between spatial and feature *K* estimates.

These tasks were specificlly designed to assess individual differences in WMc for distinct types of visual representations while controlling for perceptual-motor demands and task difficulty—factors that could otherwise confound direct comparisions of WMc-behavior associations across domains. We estimated subjects’ space- and feature-specific WMc using K-scores, defined as *K* = *S*(*H* - *F*) (Cowan, 2001), where *S* is the array size, *H* is the hit rate, and *F* is the false alarm rate. We then calculated mean *K* across three- and five-item arrays for further analyses, with a maximum possible *K* of 4. Consistent with expectation, task difficulty was well-matched at the group level: there were no significant differences in *K*-scores between the spatial and feature-based tasks among participants in Experiment 1 (Spatial: *M* = 2.74 ± 0.70 items, Feature: *M* = 2.73 ± 0.52 items, t (27) = 0.11, *p* = 0.92, *d* = 0.02), Experiment 2 (Spatial: *M* = 2.83 ± 0.69 items, Feature: *M* = 2.75 ± 0.60 items, t (27) = 0.50, *p* = 0.62, *d* = 0.10), or Experiment 3 (Spatial: *M* = 2.64 ± 0.69 items, Feature: *M* = 2.50 ± 0.75 items, t (26) = 0.88, *p* = 0.39, *d* = 0.17). The results suggest that performance was broadly comparable across the WM tasks, with no strong evidence for systematic differences in difficulty between the spatial and feature-based tasks.

Previous research shows that WMc in different visual domains is positively correlated within subjects (e.g., Eng et al., 2005), and we therefore expected some degree of covariance between spatial and feature-based scores. However, it was critical that these estimates also contained enough distinct information to allow for separate examanination of their relationship with sensorimotor learning. This criterion was met: although spatial and feature-based scores were moderately correlated in two of three experimental samples (Exp. 1: *r* = 0.34, 95% CI [-0.04, 0.63], t (26) = 1.83, *p* = 0.08; Exp. 2: *r* = 0.09, 95% CI [-0.29, 0.45], t (26) = 0.45, *p* = 0.65; Exp. 3: *r* = 0.47, 95% CI [0.11, 0.72], t (25) = 2.68, *p* = 0.01), none of the pairwise correlations exceeded 0.5, ensuring sufficient independence to estimate their unique contributions to visuomotor adaptation. Mean *K*-scores and pairwise correlations for each experiment are depicted in Fig. 1B-D. By equating memory capacity distributions across spatial and feature-based tasks in Experiments 1-3, our WM measures provided a strong foundation to test whether domain-specific or domain-general WM processes underlie individual differences in sensorimotor learning.

### Experiment 1: Does domain-specific or domain-general working memory modulate adaptation when both implicit and explicit processes are engaged?

Experiment 1 tested the association between WMc in distinct visual domains and behavioral outcomes during a standard VMA task (Fig. 2A), in which both explicit and implicit learning processes are expected to contribute—albeit to different degrees across individuals. Participants used a stylus on a touch-sensitive surface to make rapid reaching movements toward blue target circles that appeared in one of eight locations on an orthogonal display. The key manipulation occurred at the onset of the learning phase: the cursor feedback was unexpectedly rotated 45° CW or CCW relative to the actual hand position (Fig. 2A–B), thereby inducing sensorimotor errors that drive adaptation.

**Figure 2.**
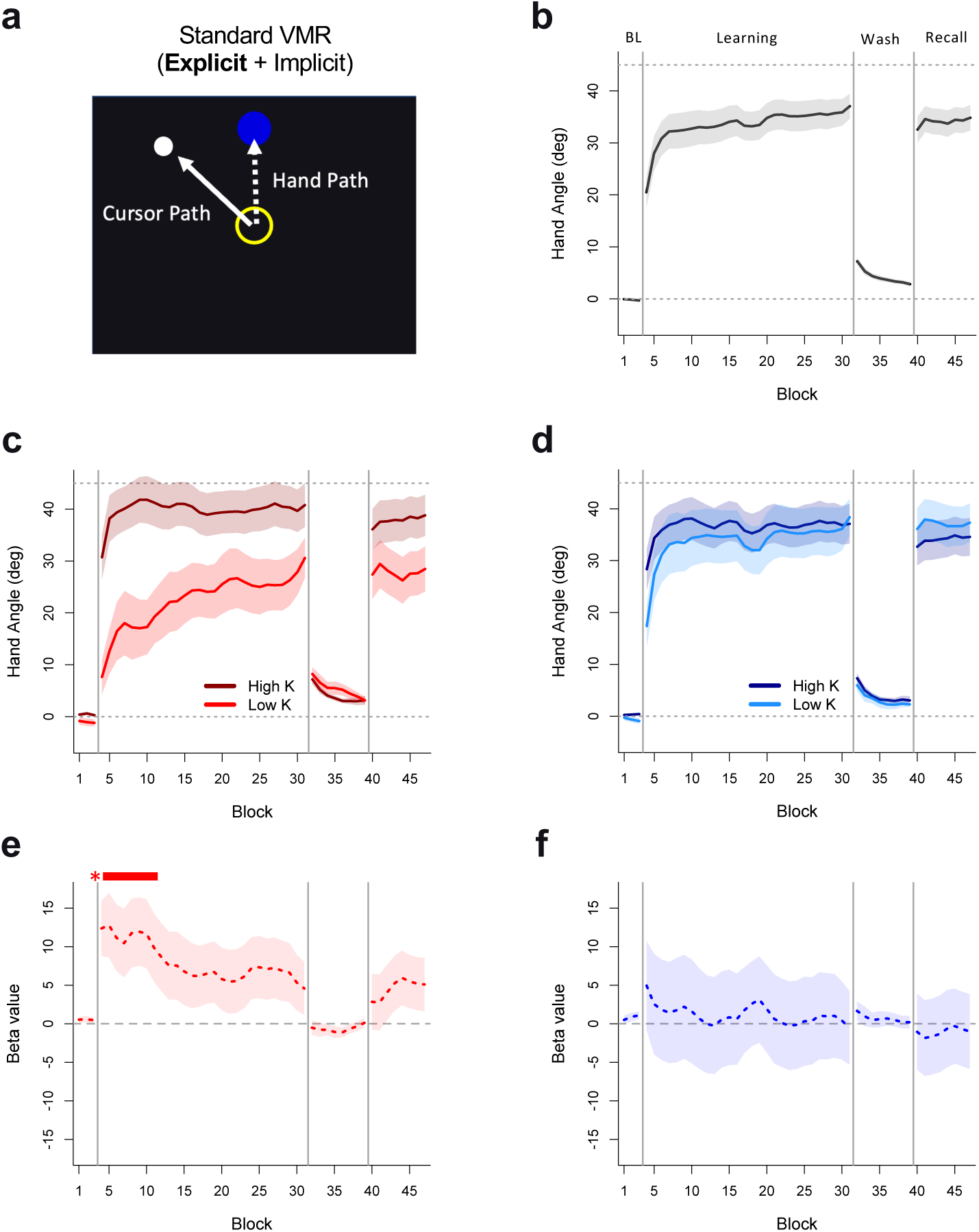
Associations between WMc and visuomotor adaptation in Experiment 1. A) Standard VMR task. During baseline and washout phases, participants received veridical cursor feedback. During learning and recall phases, continuous cursor feedback was rotated 45° relative to the hand. B) Group-averaged learning curves. Lines depict mean hand angle +/- SEM across a three-block moving average. C) Learning curves for subsets of participants with spatial WMc above the 66^th^ percentile (high K, dark red) and below the 33^rd^ percentile (low K, light red) which illustrate the interaction between spatial WMc and block during the learning phase. D) Learning curves for participants with feature WMc above the 66^th^ percentile (high K, dark blue) and below the 33^rd^ percentile (low K, light blue). E) Beta coefficients +/- SE for the continuous regression between spatial WMc and hand angle at each timepoint. The solid red bar indicates blocks with a significant positive coefficient (*p* < 0.05). F) Beta coefficients +/- SE for the continuous regression between feature WMc and hand angle at each timepoint.

Previous studies demonstrate that participants tend to leverage explicit aiming strategies to initially overcome the large, abrupt task error (Anguera et al., 2010; Holland et al., 2019; Huang & Schadmehr, 2009; Taylor & Thoroughman, 2007; 2008). As learning progresses, this strategic adjustment gives way to more implicit, SPE-driven learning, facilitiated by continuous visual feedback of the cursor position relative to the hand (Smith et al., 2006; Schadmehr et al., 2010). The magnitude of implicit learning can be estimated during the washout phase, where veridical cursor feedback is restored. This leads participants to "overshoot" in the opposite direction required to counteract the perturbation, known as an aftereffect. Finally, the perturbation is reintroduced during the recall phase to quantify the rate of relearning following previous experience.

Analysis of the resulting learning curves revealed that participants successfully adapted to the abrupt perturbation: by the end of the learning phase participants reached a mean hand angle of 37.1° (*SD* = 12.7°; Fig. 2B). Most of this change took place within the first five blocks (*M* = 28.8°, *SD* = 16.5°), a period typically dominated by explicit strategy use. During the washout phase, when veridical cursor feedback was restored, participants exhibited a significant reach aftereffect in the first block (*M* = 7.2°, *SD* = 3.2°, t (27) = 11.5, *p* < 0.001, *d* = 2.18, one sample t-test), consistent with the hypothesis that implicit learning would also contribute to adaptation.

Our primary aim was to determine whether domain-general or domain-specific visual WMc predicted learning outcomes across participants. Based on prior work (Anguera et al., 2010; 2012; Christou et al., 2016), we hypothesized that differences in spatial WMc would predict participants’ adaptation during the early stage of learning (blocks 1-5), when explicit re-aiming is expected to dominate. We also tested whether this association extended to feature-based WMc, consistent with a domain-general account. To evaluate these relationships, we employed linear mixed models with random intercepts for subjects and fixed effects for learning block, spatial WMc, feature WMc, and their interactions, with hand angle as the dependent variable. Because we hypothesized that the effects of WMc on adaptation would be dependent on task phase (e.g., only when feedback was perturbed), we fit separate models for each phase of the task (baseline, learning, washout, and recall).

Consistent with a domain-specific association between spatial WMc and adaptation, we observed a significant 3-way block × spatial × feature WMc interaction during the learning phase (*F* (1, 752) = 8.46, *p* = 0.004). Specifically, higher spatial WMc was associated with faster adaptation across learning blocks, and this benefit was stronger than any conferred by higher feature WMc. To illustrate this interaction, Fig. 2C-D depicts the learning curves associated with high (above 66^th^ percentile) versus low (below 33^rd^ percentile) WMc particiapnts separately for the spatial (red) and feature (blue) dimensions. Because the cutoff between high and low WMc is arbitraty, we created these groups only to facilitate interpretation of our primary models and did not subject them to statistical comparison. However, we computed both Cohen’s *d* and the absolute effect size for the difference in hand angle during learning for high- and low-WMc participants to contextualize the magnitude of their separation. Dividing participants based on spatial WMc yielded a large effect (Fig. 2C; *d* = 1.23; 17.4°), while the effect of feature WMc was small to negligible (Fig. 2D, *d* = 0.17; 2.8°). In contrast to the learning phase, linear models revealed no significant predictive effects of either WMc subtype on hand angle during baseline (*F* (1, 51) = .001, *p* = 0.99), washout (*F* (1, 192) = 0.20, *p* = 0.65), or recall (*F* (1, 192) = 0.15, *p* = 0.70).

While the preceding analyses establish that adaptation was differentially associated with spatial WMc during learning, we conducted follow-up analyses to more precisely identify when these effects were most pronounced. To do so, we examined continuous pairwise regressions between hand angle and WMc for each learning block (Fig. 2E-F). This analysis revealed significant associations between spatial WMc and hand angle during the first eight blocks of learning (64 trials), corresponding to the period when subjects are expected to rely primarily on explicit strategies (all *p*s < 0.05). In contrast, feature WMc had no significant association with hand angle at any point during the task. Together, these results further support a domain-specific relationship between WMc and VMA, and suggest that spatial WMc likely contributes to VMA via the selective enhancement of explicit learning.

### Experiment 2: Does spatial working memory selectively modulate explicit adaptation?

Based on the results of Experiment 1, we infer that spatial WMc selectively enhanced explicit learning processes because its effects were more pronounced during early learning. Although there is strong evidence that explicit strategies, such as re-aiming, tend to dominate during this period (Mazzoni & Krakauer, 2006; Taylor, Krakauer, & Ivry, 2014; Taylor et al., 2010; Taylor & Ivry, 2011), cumulative adaptation represents a mixture of both explicit and implicit processes, making it difficult to isolate their respective contributions. To better target the role of visual WMc in explicit learning, we modified the VMA task in Experiment 2 by removing continuous cursor feedback and instead providing only delayed endpoint feedback (Fig. 3A). This manipulation prevents online error correction and disrupts implicit learning by interferring with the processing of sensory prediction errors (Brunder et al., 2016; Izawa & Schadmehr, 2011). We predicted that if spatial WMc specifically enhances explicit learning, the domain-specific effects observed in Experiment 1 would replicate in this delayed-feedback task and persist across learning, given the diminished contribution of implicit learning.

**Figure 3.**
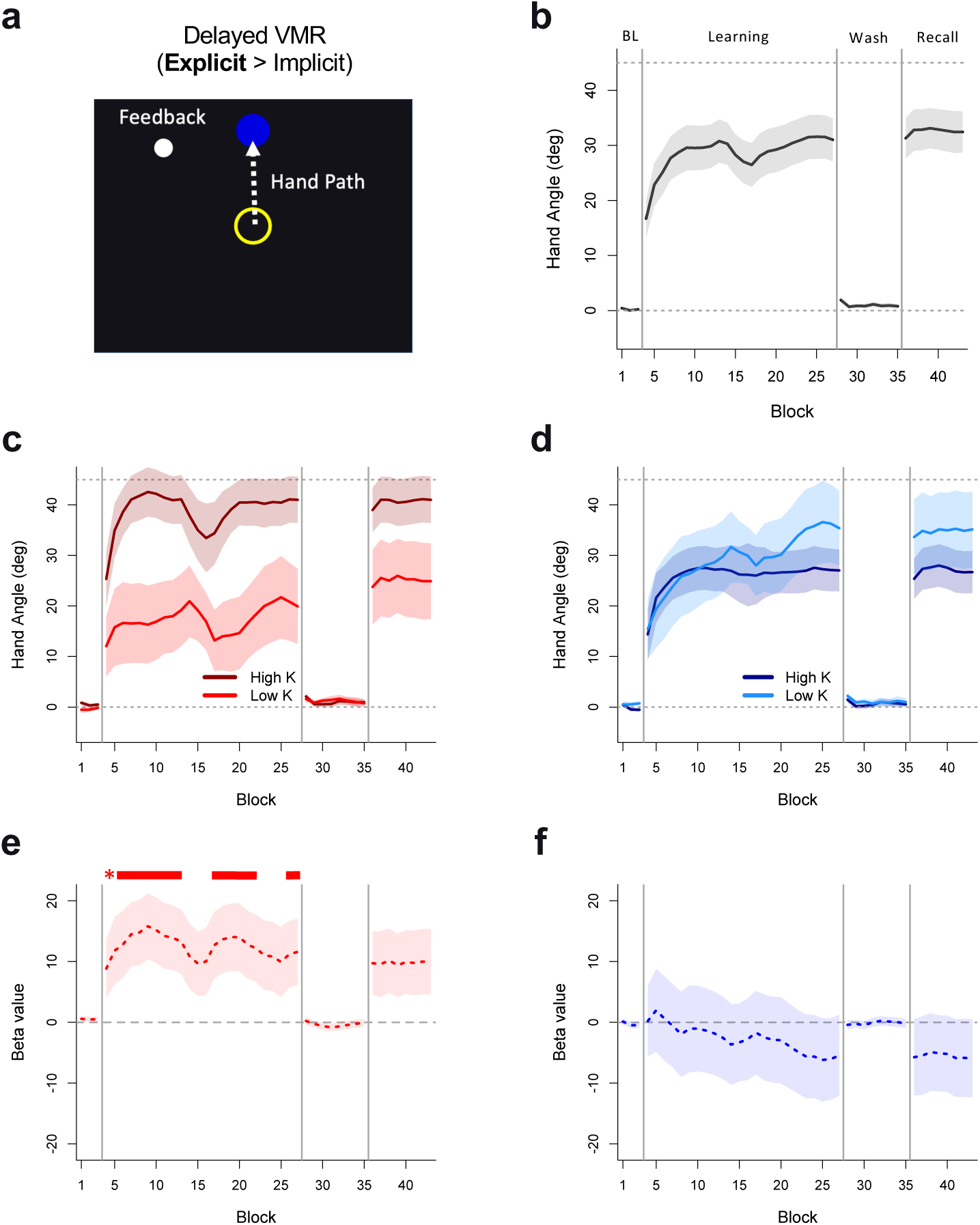
Associations between WMc and visuomotor adaptation in Experiment 2. A) Delayed VMR task. Participants did not receive cursor feedback in real time, but were provided with veridical (baseline, washout) or 45° rotated (learning, recall) endpoint feedback relative to the hand following a 500 ms delay. B) Group-averaged learning curves. Lines depict mean hand angle +/- SEM across a three-block moving average. C) Learning curves for subsets of participants with spatial WMc above the 66^th^ percentile (high K, dark red) and below the 33^rd^ percentile (low K, light red) which illustrate the interaction between spatial WMc and block during the learning phase. D) Learning curves for participants with feature WMc above the 66^th^ percentile (high K, dark blue) and below the 33^rd^ percentile (low K, light blue). E) Beta coefficients +/- SE for the continuous regression between spatial WMc and hand angle at each timepoint. The solid red bar indicates blocks with a significant positive coefficient (*p* < 0.05). F) Beta coefficients +/- SE for the continuous regression between feature WMc and hand angle at each timepoint.

In Experiment 2, participants generally learned to counteract the perturbation, reaching a mean hand angle of 24.2° (*SD* = 19.6°) during early learning (blocks 1-5) and 31.0° (*SD* = 20.6°) by the end of learning (Fig. 3B), although between-subject variability was considerably higher in the delayed sample relative to Exp. 1. Critically, the reach aftereffect was minimal during washout (*M* = 1.9°, *SD* = 2.4°, t (27) = 4.28, *p* < 0.001, *d* = 0.81, one sample t-test), suggesting that the delayed feedback manipulation successfully attenuated implicit adaptation.

By pushing participants to rely exclusively on explicit processes, we expected that spatial WMc, but not feature WMc, would exhibit a temporally sustained association with adaptation under delayed feedback. Using the same analysis approach as Experiment 1, we fit linear mixed models with fixed effects for both WMc subtype scores and learning block to test whether WMc modulated learning outcomes. Neither spatial nor feature-based WMc were associated with performance during the veridical baseline (*F* (1, 51) = 3.01, *p* = 0.09) or washout phases (*F* (1, 192) = 1.89, *p* = 0.17). As expected, WMc significantly modulated performance during the learning phase, as indicated by a significant 3-way block × spatial × feature interaction (*F* (1, 640) = 18.6, *p* < 0.001). This same 3-way interaction was also observed during the recall phase (*F* (1, 192) = 3.99 *p* = 0.047). These findings aligned with our prediction that participants with higher spatial WMc would exhibit enhanced learning relative to those with higher feature WMc when explicit strategies were emphasized (Fig. 3C-D). During learning, the difference in mean hand angle was large (21.4°) when comparing high- and low-WMc participants in the spatial domain (Fig. 3C; *d* = 1.31), in contrast to a small effect in the opposite direction (-3.3°) when dividing participants by feature-based WMc (Fig. 3D; *d* = 0.16).

To clarify when these effects emerged during learning, we examined block-specific pairwise regressions between hand angle and WMc for both memory subtypes (Fig. 3E-F). In a learning context that minimized the contribution of implicit adaptation, the benefit of higher spatial WMc on task performance both emerged early (learning blocks 2-10) and was largely sustained until the conclusion of the learning phase (blocks 14-19; 24, all *p*s < 0.05). Conversely, we found no significant association between feature-based WMc and hand angle at any timepoint.

Building on our findings from Experiment 1, these results provide further support for the hypothesis that domain-specific spatial WMc selectively enhances explicit adaptation. Moreover, we show that this association can extend well beyond the initial stages of learning when participants’ ability to implicitly refine their movements is disrupted.

### Experiment 3: Does spatial working memory capacitiy modulate implicit adaptation?

While Experiments 1 and 2 revealed a strong association between spatial WMc and learning outcomes in contexts that rely more heavily on explicit strategies, it remains unclear whether visual working memory also modulates *implicit* adaptation. One possibility is that WMc should not predict implicit adaptation at all, given that it occurs automatically and relies on dissociable neurophysiological systems from those supporting explicit strategies (Taylor et al., 2010; Izawa et al., 2012). However, an alternative hypothesis is that WMc does not exclusively support explicit processes, but instead influences the balance between explicit and implicit learning mechanisms when both are available (Albert et al., 2022; Christou et al., 2016). From this perspective, individuals with higher spatial WMc may preferentially engage explicit strategies, which in turn could suppress implicit adaptation through competitive or inhibitory interactions between systems (e.g., Boyd & Winstein, 2004; Fletcher et al., 2005; Poldrack et al., 2001).

To test these alternatives, we isolated implicit adaptation in Experiment 3 using a clamped visual feedback paradigm (Morehead et al., 2017). Participants were instructed to reach directly to targets while ignoring the cursor, which was fixed at a 45° offset relative to the target, regardless of actual hand position (Fig. 4A). Because the feedback is invariant and independent of their movement, adjusting movement based on this error signal cannot improve task performance. Nevertheless, the hand angle gradually shifted in the direction opposite the clamp, reflecting adaptation to a consistent but task-irrelevant SPE. This pattern provides a behavioral signature of implicit adaptation, as it occurs despite explicit instructions to ignore the feedback (Kim et al., 2018; Morehead et al., 2017).

**Figure 4.**
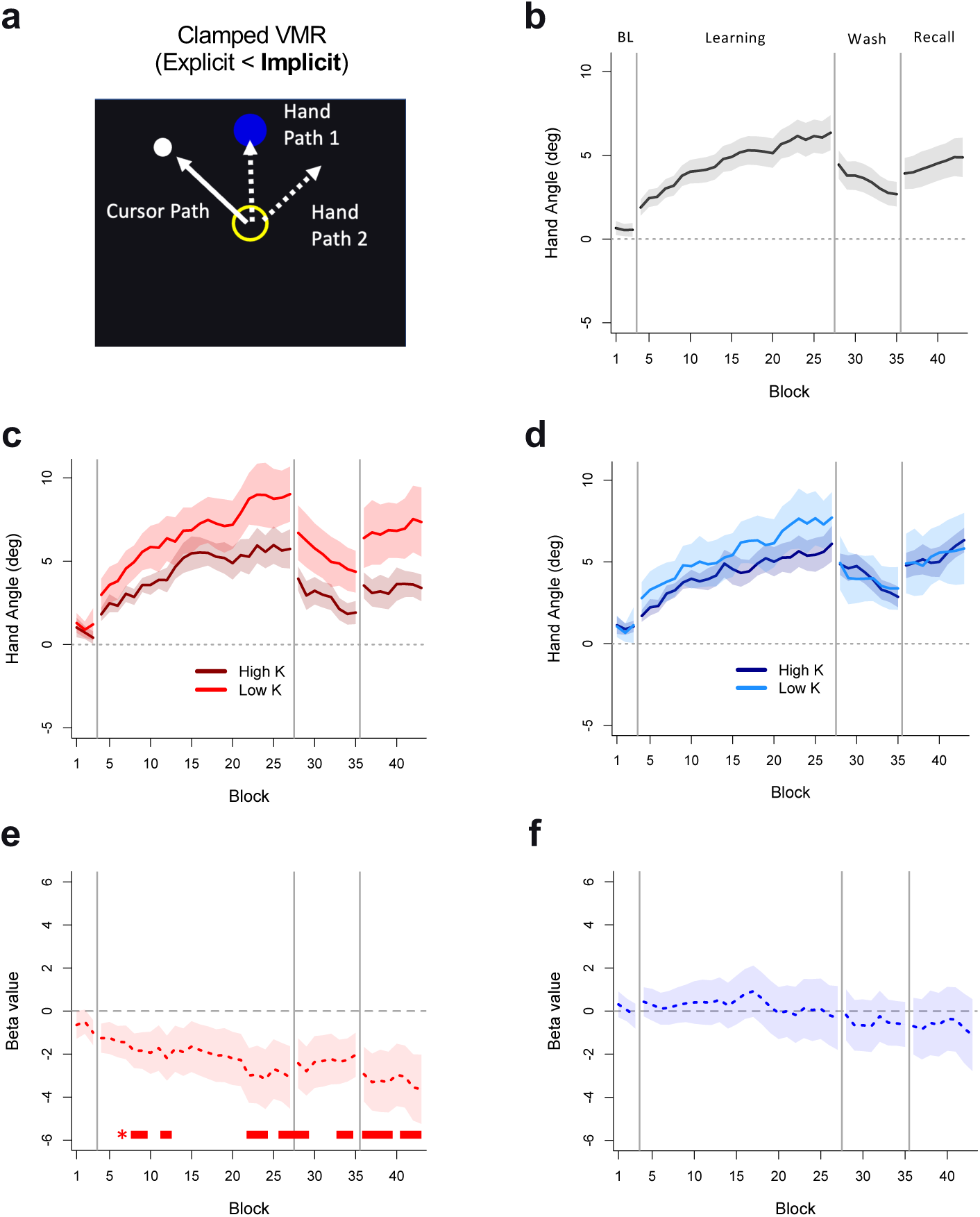
Associations between WMc and visuomotor adaptation in Experiment 3. A) Clamped VMR task. During the baseline phase, participants received continuous, veridical feedback. For learning and recall, participants were instructed to reach directly to the target and ignore cursor feedback, which was rotated 45° relative to the target and independent of hand position. No cursor feedback was provided during the washout phase for Exp. 3. B) Group-averaged learning curves. Lines depict mean hand angle +/- SEM across a three-block moving average. C) Learning curves for subsets of participants with spatial WMc above the 66^th^ percentile (high K, dark red) and below the 33^rd^ percentile (low K, light red) which illustrate the interaction between spatial WMc and block during the learning and recall phases. D) Learning curves for participants with feature WMc above the 66^th^ percentile (high K, dark blue) and below the 33^rd^ percentile (low K, light blue). E) Beta coefficients +/- SE for the continuous regression between spatial WMc and hand angle at each timepoint. The solid red bar indicates blocks with a significant negative coefficient (*p* < 0.05). F) Beta coefficients +/- SE for the continuous regression between feature WMc and hand angle at each timepoint.

As expected, the clamped feedback task led to reliable implicit adaptation, with hand angles offset by an average of 6.3° (*SD* = 5.5°) in the opposite direction of clamp by the end of learning (Fig. 4B). During the first block of washout, participants exhibited a mean reach aftereffect of 4.4° (*SD* = 4.4°, t (27) = 5.34, *p* < 0.001, *d* = 1.01, one sample t-test). Although the magnitude of adaptation and the associated aftereffect were small, we note that our design included fewer blocks and more trials per block than typical in order to match the 8 unique target locations presented in Exps. 1 and 2. Nonetheless, the observed aftereffect was proportionally larger than those observed in Experiments 1 and 2—accounting for 69.8% of final hand angle compared to 19.4% and 6.1%, respectively. This pattern is consistent with the prediction that adaptation was driven predominantly by implicit processes.

If WMc exclusively interacts with explicit processes, we would expect no differences in implicit adaptation between participants with high and low WMc under clamped feedback. Alternatively, if either domain-specific or domain-general WMc is associated with a tradeoff between explicit and implicit learning mechanisms, participants with higher memory capacity may show reduced reliance on implicit adaptation. In this case, stronger reliance on explicit learning could supress SPE-driven implicit learning, even when the task prevents strategic corrections.

To test these possibilities, we fit linear mixed models with hand angle as the dependent variable separately for each task phase, which included fixed effects for block, spatial WMc, feature WMc, and their interactions. Surprisingly, in support of the hypothesis that higher spatial WMc may indirectly suppress implicit adaptation, we observed a significant interaction between spatial WMc and block: During the learning phase, we observed a two-way interaction between spatial WMc and block (*F* (1, 617) = 8.18, *p* = 0.004), such that subjects with higher spatial WMc adapted more slowly to the clamped feedback than those with lower WMc (Fig. 4C-D). In contrast, differences in feature WMc did not modulate adaptation (*F* (1, 617) = 1.21, *p* = 0.21), although the 3-way interaction between spatial WMc, feature WMc, and learning block was also not significant (*F* (1, 617) = 2.35 *p* = 0.13).

While neither WMc subtype modulated the rate of (de-)adaptation during washout or recall, spatial WMc showed a significant main effect in both phases, (washout: *F* (1, 24) = 5.28, *p* = 0.02; recall: *F* (1, 24) = 5.48, *p* = 0.02). This suggests that group differences in adaptation observed during learning remained stable for the rest of the experiment. As expected, hand angle during baseline was not modulated by either WMc subtype (*F* (1, 47) = 0.97, *p* = 0.33). Together, these results support the interpretation that implicit adaptation was suppressed among participants with higher spatial WMc, while feature WMc had little to no independent effect. Despite the positive correlation between spatial and feature WMc estimates among subjcets in Experiment 3 (see Fig. 1D), the effect sizes for the difference in hand angle between high and low*-K* subjects depicted in Fig. 4C-D reveal a medium effect of spatial WMc (*d* = 0.71; -2.2°) compared to a small effect of feature WMc (*d* = 0.31, -1.3°). Importantly, only the spatial WMc effects were statistically significant in the linear mixed models.

To further examine the time course of the observed suppression, we conducted pairwise block-level regressions between hand angle and spatial WMc (Fig. 4E-F). This analysis indicated significant negative associations at multiple time points during learning (blocks 5-6, 9, 19-21, 23), washout (blocks 1-2, 6-7), and recall (blocks 1-4, 6-8). The relationship between spatial WMc and reduced implicit adaptation became more prounounced with increasing time on task, suggesting a cumulative or sustained suppression effect. Once again, no significant association was observed between feature WMc and hand angle at any time point.

To our knowledge, these results are the first to link individual differences in cognitive capacities among healthy subjects to implicit adaptation. Even in the absence of meaningful explicit strategies, participants with higher spatial WMc were less sensitive to the noncontingent visual feedback than those with lower WMc. Moreover, the findings from Experiment 3 reinforce the conclusion that the modulatory effects of visual working memory on VMA are specific to the spatial domain and are therefore unlikely to reflect differences in domain-general WMc.

### Domain-specific Spatial WMc Drives Model Explanatory Power Across Three Types of Visuomotor Adaptation

Across three independent samples and task manipulations, our analyses suggest that spatial WMc selectively modulates sensorimotor learning outcomes—facilitating explicit learning (Experiments 1 and 2) and suppressing implicit learning (Experiment 3). However, because our primary statistical models included fixed effects for learning block, spatial WMc, and feature WMc simultaneously, we conducted stepwise regression analyses to clarify the individual explanatory contributions of each WMc subtype. Specifically, we contrasted the marginal R^2^ (variance explained by fixed effects alone; Nakagawa & Schielzeth, 2013) associated with a null (block-only) and full (block × spatial × feature) model with intermediate models that included either spatial or feature WMc alone as fixed effects. For all tested models, the dependent variable was blockwise hand angle during the learning phase.

In all three experiments, spatial WMc accounted for the vast majority of fixed-effect variance explained by the full model (Exp. 1: R^2^ = 0.141, 97.9%; Exp. 2: R^2^ = 0.167, 94.4%; Exp. 3: R^2^ = 0.184, 68.9%), whereas models including only a fixed effect for feature WMc explained litte to no variance above the null model (Fig. 5). Notably, despite the significant positive correlation between WMc subtypes in Experiment 3 (Fig 1D), feature WMc provided no explanatory value for implicit adaptation—underscoring the unique predictive value of spatial WMc. Indeed, even in a purely implicit context, spatial WMc was a more powerful predictor of participants’ adjustments in hand angle than time on task. Taken together, these results bolster the conclusion that spatial WMc plays a distinct role in shaping sensorimotor learning and demonstrates that this relationship robustly generalizes across a range of VMA contexts.

**Figure 5.**
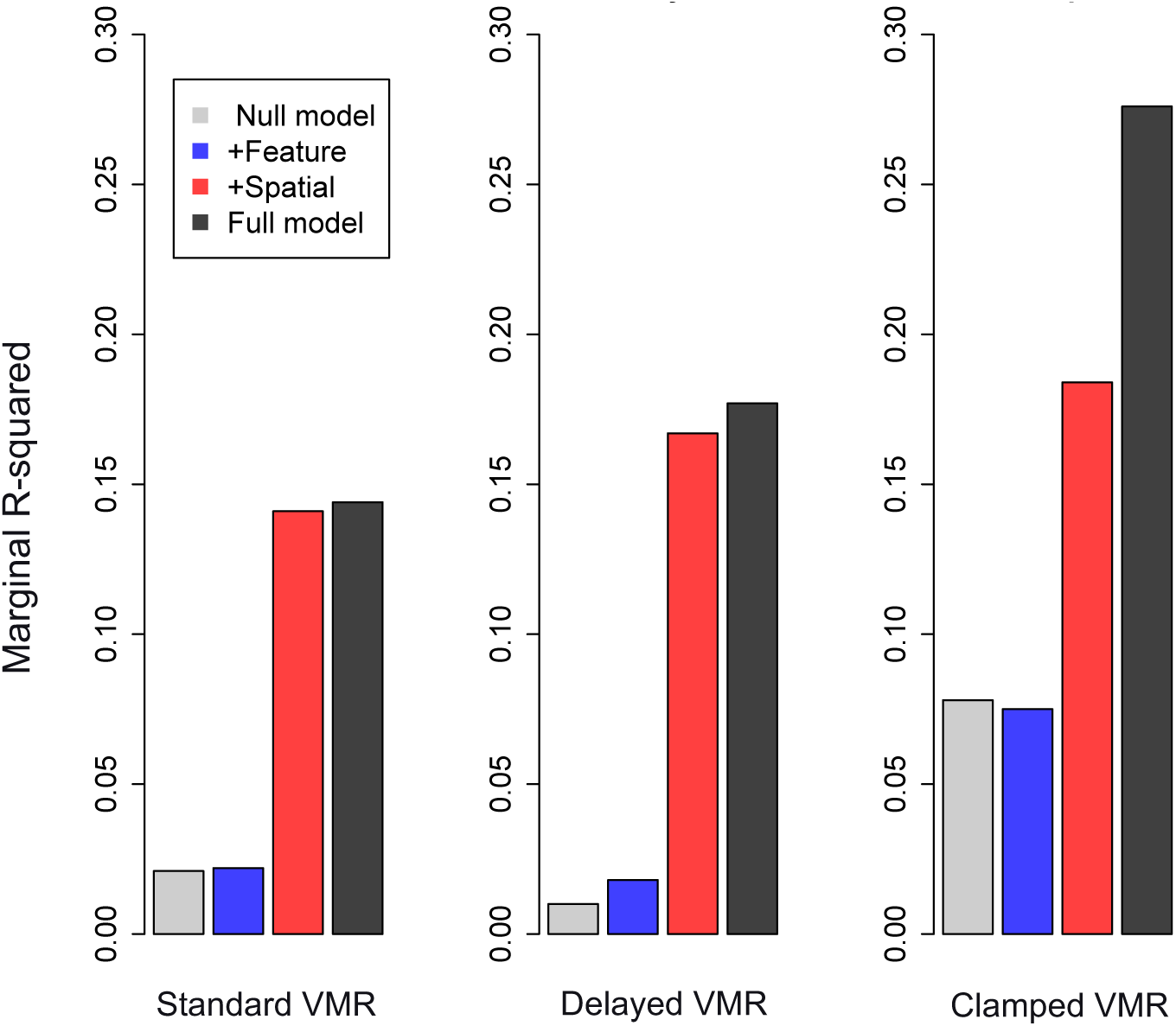
Variance explained by fixed effects across experiments. Bars depict the marginal R^2^ for linear mixed models predicting hand angle during each respective learning phase with fixed effects for block only (null model, gray), block and feature WMc (blue), block and spatial WMc (red), and a full model including all fixed effects (black).

## Discussion

Across three VMA tasks that differentially engage explicit and implicit learning mechanisms, we found strong, convergent evidence that domain-specific spatial WM, but not feature-based WM, modulates individual differences in adaptation. In tasks that involved explicit adaptation, higher spatial WMc consistently predicted greater performance, while feature-based WMc showed no reliable association with performance outcomes. Unexpectedly, we also observed significant individual variability in implicit adaptation that could be explained spatial WMc: participants with higher *K*-scores showed suppressed adaptation to clamped feedback. Together, these results clarify the format of WM representations that support visuomotor adaptation and raise new questions about how explicit and implicit learning systems interact—especially in contexts where both are potentially operative.

Our findings build on and critically extend previous research implicating visuospatial working memory with explicit components of VMA (Anguera et al., 2010; 2012; Christou et al., 2016; Holland et al., 2019). For example, Anguera and colleagues (2010) reported that performance on a 2D card rotation task—thought to recruit visuospatial WM—correlated with faster adaptation during early learning in a VMA task. The authors inferred that WM supports explicit adaptation, as this association was restricted to early learning, a phase known to be selectively disrupted by dual task interference (Eversheim & Bock, 2001; Taylor & Throroughman, 2007; 2008). In Experiment 1, by modeling this association across blocks, we confirmed that spatial WMc selectively modulated adaptation in early learning. However, it is important to note that both our study and prior work assume that explicit contributions are elevated during early learning. This assumption is supported by past studies, but the transition between explicit and implicit learning is not discrete and neither process can be isolated in standard tasks.

We designed Experiment 2 to provide a more stringent test of whether WMc specifically supports explicit adaptation by introducing delayed feedback (Brunder et al., 2016), a manipulation known to disrupt implicit learning. In this context, we hypothesized that participants would need to rely on explicit strategies throughout the entire learning phase, not just during the early stages. Therefore, if spatial WMc facilitates explicit learning, its influence should persist across learning in the delayed feedback pardigm. This prediction was confirmed, suggesting that spatial WMc supports early learning in standard VMA tasks precisely because explicit strategies are the predominant contributor. When implicit adaptation is disrupted, spatial WMc continues to explain the individual differences in adaptation. This observation underscores the importance of domain-specific spatial WM resources in motor learning contexts where implicit adaptation is impaired or unavailable, such as in patients with cerebellar disorders (e.g., McDougle et al., 2022; Wong et al., 2019).

Previous work has proposed several mechanisms to explain the observed relationship between spatial WMc and explicit adaptation. McDougle and Taylor (2019) described two dissociable strategies, parametric rotation and response caching, that are recruited in a context-dependent manner to enable people to overcome visuomotor rotations. Their work suggests that the optimal strategy depends critically on the number of possible target locations. When the number of target locations exceeds the capacity of visual WM (typically 3-4 items), participants rely on parametric rotation. This strategy is inefficient but not limited by WM capacity, and its use is typically indexed by RTs that scale with the magnitude of rotation, mirroring effects seen in mental rotation tasks. In contrast, when the number of targets locations is within the capacity limits of WM, participants exhibit uniformly fast RTs that do not vary with rotation magnitude. This pattern is thought to reflect response caching, where the action plan for each target is stored and retrieved directly from memory.

Critically, due to the structure of our tasks, the finding that higher spatial WMc facilitates explicit adaptation is incompatible with a response caching mechanism. In all experiments, targets were randomly selected from eight possible locations—well beyond the capacity limit of visual WM (3-4 items). Consequently, participants were unlikely to have stored discrete action plans for each target location. Between the two proposed strategies, these findings more strongly support the use of parametric rotation in Experiments 1 and 2. If so, our results suggest that spatial WM—even in the absence of rotational demands—may underlie people’s ability to implement and maintain a parametric strategy. However, this relationship may depend on how spatial WM is assessed. Tasks that involve rotational transformations (e.g., Angeura et al., 2010) may track strategic adjustments to rotational perturbations in a different manner than the task employed in our experiments.

In contrast to Experiments 1 and 2, where explicit strategies were expected to partially or primarily drive adaptation, Experiment 3 isolated implicit adaptation driven using clamped feedback, which elicits learning driven by sensory prediction error (Kim et al., 2018; Morehead et al., 2017). In this context, a plausible expectation was that that neither spatial nor feature-based WMc would predict behavioral outcomes, given that implicit adaptation is automatic and operates largely outside conscious awareness. Moreover, because participants were instructed to ignore the noncontingent feedback, WM-dependent strategies were ostensibly irrelevant. We therefore treated the clamped task as a control condition to strengthen inferences about the role of WMc in explicit adaptation. As intended, this approach yielded larger relative aftereffects than those observed in Experiments 1 and 2, consistent with the goal of driving implicit recalibration, which cannot be readily suppresed. Surprisingly, however, spatial WMc selectively *suppressed* adaptation such that individuals with higher spatial WMc were less responsive to the clamped feedback. This effect emerged almost immediately and became increasingly prounounced over time, suggesting that individual differences in spatial WM can shape not only explicit strategy use, but also the operation of implicit learning mechanisms under certain conditions.

The suppression of implicit adaptation among individuals with higher spatial WMc aligns with the idea that WM abilities governs tradeoffs between explicit and implicit learning systems in a “push-pull” manner (Christou et al., 2016, Holland et al., 2019). That is, individuals with high spatial WMc may preferentially engage explicit re-aiming strategies, reducing reliance on the implicit system, whereas those with lower WMc may rely more heavily on implicit recalibration to compensate for limited strategic flexibility. This competition between systems is well documented in standard VMA tasks, where increased engagement of explicit strategies often correlates with attenuated SPE-driven learning (Albert et al., 2022; Benson et al., 2011; Christou et al., 2016; Fernandez-Ruiz et al., 2011).

Critically, however, the current results are incompatible with this competition account. Unlike standard adaptation tasks that allow for strategy use, clamped feedback eliminates the relevance of explicit strategies. Under such conditions, no competition between learning systems should arise—yet spatial WMc still predicted reduced implicit adaptation. This unexpected finding suggests a more obligatory or structural interaction between learing systems not anticipated by existing models (Albert et al., 2022; Heald et al., 2021). At a minimum, these findings demonstrate that the implicit learning system is not functionally isolated from higher-level cognitive capacities, such as working memory. Put differently, individual differences in spatial working memory—and potentially other latent cognitive capacities—may shape the balance between implicit and explicit learning systems, even in contexts where only one system is engaged.

From an active inference perspective, spatial WMc may reflect variations in the precision weighting assigned to top-down predictions in sensorimotor control (Friston, 2010; Friston et al., 2015). Specifically, individuals with higher spatial WMc may more confidently engage explicit, strategy-based learning processes, effectively down-weighting incoming signals that would otherwise drive implicit recalibration (cf. Daw, Niv, & Dayan, 2005). In the context of sensorimotor adaptation, greater reliance on explicit strategies could reduce the relative contribution of implicit learning—particularly if the system deems the top-down representations sufficiently precise (Taylor, Krakauer, & Ivry, 2014). Critically, higher spatial WMc would not merely reflect a capacity to remember target locations, but also an enhanced ability to attend to and encode visual feedback with high fidelity—amplifying task-relevant cues while filtering out task-irrelevant or noisy signals.

However, under normal conditions, task error (TE) and SPE are conflated in the same feedback signal, raising the question of how spatial WM might selectively amplify one signal over the other. Within an active-inference framework, the stable internal representations afforded by robust spatial WM guide top-down attention to the discrepancies most relevant for explicit, goal-directed corrections—thereby differentially weighting these error signals. Even though implicit recalibration remains continuously operative (Mazzoni & Krakauer, 2006), higher spatial WM may effectively upweight TE, reducing the relative impact of the slower, implicit recalibration process driven by SPE. Consequently, there is no need to disentangle TE and SPE; rather, attentional and WM-based capacities skew the system’s confidence in favor of TE, thereby increasing reliance on explicit strategies and resulting in greater explicit adaptation overall.

Our current results build on research suggesting that visual WM is supported by domain-specific subsystems or “buffers” dedicated to processing different kinds of sensory information (Baddeley et al., 1999; Baddeley, 2003; Logie, 1995) and that these distinctions are highly relevant in the study of individual differences (Nozari & Martin, 2024; Zimmer, 2008). Numerous studies have revealed behavioral and neuropsychological dissociations between spatial and feature-based WM via selective interference effects (e.g., Della Salla, 1999; Goodale & Milner, 1992; Goodale, Westwood, & Milner, 2004; Hyun & Luck, 2007; Logie & Marchetti, 1991). Despite the fact that spatial and feature-based WMc were positively correlated, we found that performance in the spatial task exclusively predicted learning outcomes in each context. On one hand, the shared variance between these measures may reflect domain-general executive control processes that contribute to WM regardless of the sensory representations involved (Courtney, 2004; Serences et al., 2004). However, our findings suggest that differences in general executive function are not solely responsible for driving individual differences in VMA. Instead, the storage of spatial information in particular appears to play a unique role in support of explicit adaptation, highlighted by the substantial variance explained by spatial *K* in our model comparisons (see Fig. 5).

The domain-specificity of visual WM is thought to reflect the divergence of dorsal (“where”) and ventral (“what”) visual pathways for the representation of spatial coordinates and object features, respectively (Goodale & Milner, 1992; Goodale, Westwood, & Milner, 2004). Thus, the selective effects of spatial WMc on VMA indirectly point to an involvement of dorsal stream structures such as the intraparietal sulcus (IPS) and frontal eye fields (FEF), which are known to support both spatial WM and visually guided action. Neuroimaging studies have consistently demonstrated the engagement of parietal cortex in mental rotation tasks, such that its activity scales with the magnitude of rotation but not visual input itself (Cohen et al., 1996; Heil, 2002; Jordan et al., 2001; Suchan et al., 2006). IPS and FEF are retinotopically organized, underscoring their contributions to spatial attention (Grefkes & Fink, 2005; Scolari et al., 2015) and spatial WM (Curtis & D’Esposito, 2003; Curtis et al., 2004). For example, using a delayed saccade task where a spatial cue would be consistent or inconsistent with the location of a subsequent eye movement, Curtis and colleagues (2004) showed that these regions support dissociable WM codes that enable goal-directed action: During memory delays, the IPS represents previously encoded spatial information, where FEF represents prospective spatial information to guide future eye movements. Accordingly, one question for future research is whether spatial WM processes in VMA specifically support a retrospective mechanism (e.g., encoding previous errors), prospective mechanism (e.g., target-dependent motor planning), or both.

More work is needed to clarify the precise nature of the spatial representations involved in VMA. The current study employed a passive spatial WM task which required participants to encode and maintain discrete stimulus locations (e.g., Christou et al., 2016). This task was well-suited for our purposes because it enabled a direct comparison to memory for the same stimuli on an orthogonal dimension (color). However, other studies have implicated mental rotation in VMA (Anguera et al., 2010; Guo & Song, 2023; Holland et al., 2019). Although mental rotation likely recruits spatial WM, these processes are not the same: while rotation requires a continuous mental transformation by definition, discrete spatial WM does not. This distinction is highly relevant for sensorimotor learning. Cerebellar patients, who exhibit VMA deficits stemming from an inability to integrate sensorimotor prediction errors (Smith & Shadmehr, 2005; Tseng et al., 2007), also exhibit impaired performance in other cognitive tasks requiring continuous mental transformations (McDougle et al., 2022). Interestingly, these same patients display relatively normal WM performance in discrete spatial tasks (McDougle et al., 2022) and may even compensate for motor learning impairments by relying more heavily on spatial WM resources (Wong et al., 2019). Thus, mental rotation tasks may probe additional, cerebellum-dependent processes which track VMA performance in qualitatively different ways than observed here (see also Anguera et al., 2010; Holland et al., 2019).

In summary, across three experiments we demonstrate that individual differences in sensorimotor learning are modulated by spatial working memory capacity. These results underscore the domain-specific organization of visual working memory the pervasive interaction between cognitive abilities and adaptive motor control.

## Methods

### Participants

A total of 90 healthy adult volunteers (59 female, 30 male, 1 unreported; *M*_age_ = 20.9, *SD* = 4.4) from the Brown University community participated for course credit or $10/hour across the three experiments (*n* = 30 for each group). The desired sample size of 30 subjects per experimental group was determined based on the sample sizes for comparable visuomotor adaptation studies from our lab and elsewhere (Brunder et al., 2016; Im, Liddy, & Song, 2022; Morehead et al., 2017, Wang et al., 2022; Song & Bédard, 2015), which yielded large effect sizes (𝜂_p_^2^ > 0.26) for primary learning outcomes with group sample sizes of 20 or fewer; we recruited 10 additional subjects per group to support individual difference analyses related to WMc. All participants were right-handed, reported normal or corrected-to-normal vision and normal color vision, were free of neuromuscular impairment, and were naive to the visuomotor rotation task.

For Experiment 1, we excluded one subject due to technical failures and another subject due to extreme baseline reach trajectories (>3 *SD* from the mean). For Experiment 2, we excluded two subjects from analyses for (1) failure to make straight reaching movements on >20% of learning trials and (2) not adhering to task instructions. Finally, two subjects were removed from analyses for Experiment 3 due to technical failures resulting in a loss of data. One subject in Experiment 3 exhibited chance-level performance on the feature WM task, leading to the exclusion of their feature *K* estimate from analyses. After exclusions, the final analyzed sample size was *n* = 28 for each experimental group. All protocols were approved by the Brown University Institutional Review Board, and participants provided written informed consent in accordance with the Declaration of Helsinki.

### Apparatus

For Experiment 1, participants were seated in a dimly lit room at a distance of 90 cm from a Viewsonic G90fB CRT monitor measuring 36 x 24 cm (resolution: 1024 x 768). For Experiments 2 and 3, participants were seated 65 cm from a Viewsonic PJD6221 projector screen measuring 45 x 33.8 cm (resolution: 1024 x 768). All other aspects of the visual displays and apparatus were identical across experiments unless otherwise noted.

Holding a stylus in their right hand, participants made reaching movements by sliding the stylus tip across a digitizing tablet (Magic Touch; Tyco Touch, Inc.) placed flat on a table in front of them, which was aligned with the participant’s midsection and the center of the monitor. Visibility of the right arm and hand was occluded by a rectangular section of black opaque fabric positioned underneath the projector screen and 20 cm above the touch surface. Visual stimuli were created using MATLAB 2016b and presented via Psychophysics Toolbox (v3, Kleiner et al., 2007) on a PC with a 60 Hz refresh rate.

We used the same apparatus and displays as detailed above to administer the spatial and feature-based working memory tasks for each respecitve experiment, with the exception of placing a keyboard in front of participants to record responses in lieu of the touch surface.

### Experimental Design

#### Working Memory Tasks

To estimate working memory capacities, participants from each experimental group completed identical change-detection tasks which required them to remember the locations (spatial task) or colors (feature task) of dot arrays over a brief delay period (Fig. 1A). The working memory tasks were always administered before the visuomotor rotation task, and at this time subjects were naive to the second phase of the experiment (VMR). All subjects completed two 30-trial blocks of each WM task (120 trials in total), and the task blocks were presented in an ABBA order which was counterbalanced across participants. Memory load was randomized across trials within each block such that 50% of trials presented a 3-item array (low load) and 50% of trials presented a 5-item array (high load) at encoding. For exploratory analysis, participants completed 30 additional trials of the spatial and feature tasks with 7-item arrays; these trials were not used to compute *K* because they exceed the theoretical limit of working memory capacity.

Before each task block, participants were explicitly instructed to attend to the relevant stimulus dimension (dot location or dot color) and that the other dimension would not be relevant for their responses. Procedurally, each trial began with a white fixation point (0.4° diameter) presented on a black background for 2 s, which also served as the interstimulus interval. Following fixation, for the enconding period an array of three or five uniquely colored dots (0.6° diameter) were presented in a seemingly random spatial arrangement. This was followed by a 2.5 s fixation-only delay period. Finally, one colored dot was presented for 2 s, and during this time participants were required to make a keypress corresponding to whether the dot was a match or non-match with one of the locations (spatial task) or one of the colors (feature task) seen in the initial encoding display. Participants received auditory feedback in the form of a high-pitched (correct) or low-pitched (incorrect) beep on each trial when a response was made.

We conducted pilot studies with indepdendent participants prior to designing the final WM tasks to calibrate task difficulty, such that performance (estimated *K*) on the spatial and feature tasks would be approximately matched. For the spatial task, our final design drew dot locations randomly from 16 possible spatial positions. These positions corresponded to two concentric rings, with 8 positions within each ring. The "outer" ring positions were located 4° from fixation and were separated by 45° clockwise, starting at 12 o’clock relative to central fixation. The "inner" ring positions were located 2.6° from fixation, and their possible clockwise positions were offset by 22.5° relative to those of the outer ring.

To associate each dot with a distinct visual feature, we drew dot colors randomly from a list of 8 possible colors. We selected colors to maximize both psychological and perceptual separability (e.g., Boynton, 1989) for the feature-based task, including red (RGB: 255, 0, 0), blue (0, 255, 0), green (0, 0, 255), yellow (255, 255, 0), pink (255, 0, 255), cyan (0, 255, 255), maroon (160, 0, 0), and orange (255, 160, 0).

Thus, on each trial for both WM tasks, the dot positions and colors were drawn randomly from the same list of 16 possible locations and 8 possible colors. We also applied the same probabilities for stimulus change (match/non-match) to the single dot presented for retrieval across tasks. In 50% of all trials, the dot position changed to a random location that was not presented in the encoding display, and in 50% of trials, the dot color changed to a random color not seen in the encoding display from the remaining possibilities. Thus, changes (or a match) on one stimulus dimension were completely orthogonal to changes on the other dimension. The two tasks differed only with respect to which stimulus dimension needed to be attended to achieve correct responses.

### Experiment 1: Standard VMR

Each trial began with participants moving the cursor (white circle, 0.25 cm diameter) to a starting position located in the center of the screen, indicated by a yellow annulus (1 cm diameter). Before target onset, participants held the cursor in the starting position for a minimum of 1 s, plus a random interval between 0 - 1 s such that the timing of target onset was not completely predictable. Following this interval, the target (blue circle, 1 cm diameter) appeared in one of eight possible locations. Targets were located 6.75 cm from the center of the screen and each possible location was separated by 45°, beginning at 12 o’clock relative to the starting position. Each target location was encountered in a pseudorandom order across blocks of 8 trials.

Participants were instructed to make a rapid, straight-line movement to slice through the target with the cursor. We encouraged participants to overshoot the target and discouraged them from making any corrective movements during a trial. Both the target and cursor remained visible for the full trial duration (3 s). When the cursor passed the distance of the target (6.75 cm), participants were provided with stationary feedback about their movement error relative to the target for the remainder of the trial. In between trials, cursor position was masked and replaced by a distance circle which guided participants back to the starting position.

There were four task phases: (1) baseline (40 veridical trials), (2) learning (240 rotation trials), (3) washout (80 veridical trials), and (4) recall (80 rotation trials). The baseline phase served as a control to assess whether WMc was associated with any biases in reach direction not attributable to learning. In the primary learning phase, continuous cursor feedback was abruptly rotated either 45° CW or CCW relative to the hand position, where the direction of this rotation was held constant within participants (but counterbalanced across participants). In the washout phase, participants received continuous, veridical feedback about their hand position, and were expected to de-adapt to the rotation. Finally, during the recall phase, the same rotation participants encountered during learning was abruptly reintroduced. A short (<30 s) break was provided between each experimental phase and at the midpoint of the learning phase.

### Experiment 2: Delayed VMR

Unless otherwise noted, all aspects of the visuomotor rotation task and display parameters were identical to those of Experiment 1. For Experiment 2, the key manipulation and departure from the standard VMR design was the introduction of delayed cursor feedback in order to limit implicit adaptation and emphasize explicit rotation strategies (Brunder et al., 2016). Participants again initiated trials by moving the cursor into the starting position, during which time the cursor remained visible. After the onset of the target, the cursor disappeared, and participants made a rapid movement to slice through the target in absence of continuous cursor feedback. Finally, 500 ms after the (invisible) cursor passed the 6.75 cm target distance, we provided fixed endpoint feedback about the location of the cursor relative to the target for the remainder of the trial. The total trial duration was extended to 3.5 s (relative to 3 s in Exp. 1) to account for the additional 500 ms delay. We used the same task phases as Exp. 1 with the exception of 36 fewer learning trials: (1) baseline (40 veridical trials), (2) learning (208 rotation trials), (3) washout (80 veridical trials), and (4) recall (80 rotation trials).

### Experiment 3: Clamped VMR

Unless otherwise noted, all aspects of the visuomotor rotation task and display parameters were identical to those of Experiment 1. The key manipulation for Experiment 3 was the incorporation of clamped visual feedback (Morehead et al., 2017) to isolate implicit adaptation. Critically, for the clamped feedback task we explicitly instructed participants to ignore the cursor feedback and reach directly to the target. In contrast to Exp. 1, where continuous cursor feedback was rotated 45° relative to the hand, during learning and recall phases in Exp. 3 cursor feedback was rotated at a fixed offset of 45° CW/CCW relative to the position of the target. Thus, while the cursor feedback matched the timing components of the hand movement (reaction time and velocity), the angular offset of the feedback was entirely independent of the hand movement. Participants completed four task phases: (1) baseline (40 veridical trials), (2) learning (208 clamped rotation trials), (3) washout (80 no-feedback trials), and (4) recall (80 clamped rotation trials). Note that for the washout phase in Exp. 3, no cursor feedback was provided to obtain a purer measure of the reach aftereffect indicative of implicit adaptation.

### Reach Analyses

Reach analyses were performed in MATLAB R2023b and were the same across Experiments 1–3. Consistent with the procedures of our previous studies (e.g., Im, Liddy, & Song, 2022; Wang et al., 2022; Song & Bédard, 2015) movement analyses were restricted to the center-out movement to the target. Hand position coordinates were filtered using a second-order, low-pass Butterworth filter with a 6-Hz cutoff. Hand position data were transformed to a common reference frame with the target located at the 12 o’clock position. Hand displacements were obtained using a first-order backward difference. Net hand displacements were computed as the square root of the sum of the squared x- and y-displacements. Hand tangential velocity was obtained by dividing the net hand displacements by the sampling interval (1/200 Hz = 0.005 s). Movement onset and offset were identified when the hand tangential velocity first exceeded and fell below 5% of peak velocity. Each movement trajectory and velocity profile were visually inspected to ensure that the entire movement was captured by the movement detection algorithms. Hand angle, the angular difference between the target and the initial movement direction, was used to measure performance in the VMA tasks. The initial movement direction was defined as the line connecting the hand position at movement onset and peak velocity in each trial. A positive hand angle denoted clockwise deviations relative to the target, whereas a negative hand angle denoted CCW deviations. Trials were excluded if the hand angle exceeded ±90°, indicating a movement in the wrong direction, or deviated more than 25° from the median of the five previous and successive trials. In Experiment 1, the total number of trials removed were 445 (3.6%) of 12,320 trials. In Experiment 2, the total number of trials removed were 532 (4.7%) of 11,424 trials. In Experiment 3, the total number of trials removed were 246 (2.2%) of 11,424 trials. Adjusting the outlier removal parameters within reasonable limits did not affect the reported results. Hand angle was averaged across mini-blocks of eight successive trials, one per target location. Finally, we temporally smoothed the data for analysis and plotting by computing a 3-block (24 trial) moving average of hand angle across mini-blocks. We note that this temporal smoothing procedure did not lead to any qualitative changes in results in comparison to the unsmoothed data.

### Statistical Analyses

Statistical analyses for working memory data and processed reach data were carried out in R (version 4.3.1; https://www.r-project.org/). Primary statistical effects were evaluated via linear mixed models implemented in the “lme4” R package using restricted maximum likelihood estimation. For all experiments and task phases, the models included continuous fixed effects for block, spatial WMc (*K*), feature WMc (*K*), and their interactions, with subject as a random effect and hand angle as the dependent variable. Model residuals were visually inspected for non- linearity, heteroscedasticity, and outliers. Reported *r*-values for the linear correlations between spatial and feature-based WMc estimates reflect Pearson’s *r*. Post-hoc comparisons were computed via paired, one-sample, or two-sample t-tests, where appropriate. Beta coefficients and standard errors for the block-level associations between WMc and hand angle were computed using simple linear regression. All statistical comparisons were two-tailed. Reported standardized effect sizes reflect Cohen’s *d* for one- and two-sample t-tests, *d_z_* (*M*_diff_ / *SD*_diff_) for paired t-tests, and marginal R-squared for linear mixed models (see Fig. 5; Nakagawa & Schielzeth, 2013).

## Data Availability Statement

Raw and preprocessed data will be shared upon request during peer review at the Open Science Framework https://osf.io/ntgs7/ and will be made publicly available upon acceptance at https://osf.io/ntgs7/.

## Code Availability

Experiment code and analysis code necessary to replicate these results will be shared upon request during peer review at the Open Science Framework https://osf.io/ntgs7/ and will be made publicly available upon acceptance at https://osf.io/ntgs7/.

## Acknowledgements

This work was supported by NSF BCS-2341363 and NSF BCS-2043328 to J.H.S.

We thank Alex Daskalopoulos and Anel Zhussubali for their assistance with data collection.

## Author Contributions

S.R.O. – Conceptualization, Data Curation, Formal Analysis, Investigation, Methodology, Project Administration, Software, Validation, Visualization, Writing - Original Draft Preparation. J.L. – Conceptualization, Data Curation, Methodology, Software, Writing - Review & Editing. J-H.S. – Conceptualization, Funding Acquisition, Methodology, Project Administration, Resources, Supervision, Writing - Review & Editing.

## Notes

### Competing Interest Statement

The authors have declared no competing interest.

### Summary of Updates

Added data availability and code availability statements; included additional details in the Methods; revised Figure 3 (recall phase); revised Figure 4D and 4F to reflect exclusion of one subject.

